## Supplementary Materials for "Regional Topological Aberrances of White Matter- and Gray Matter-based Functional Networks for Attention Processing may Foster Traumatic Brain Injury-Related Attention Deficits in Adults"

**Table S1.** Nodes for White Matter Functional Brain Network Construction

| **Anatomical Regions** | **Abbreviations** | **Local Maxima in MNI coordinates of power spectrum** | | |
| --- | --- | --- | --- | --- |
|  |  | **x** | **y** | **z** |
| Genu_of_corpus_callosum | GCC | -2 | 24 | 10 |
| Body_of_corpus_callosum | BCC | 2 | -28 | 22 |
| Splenium_of_corpus_callosum | SCC | 0 | -38 | 12 |
| Fornix | FX | 0 | -8 | 18 |
| Cerebral_peduncle_L | CP_L | -16 | -12 | -6 |
| Anterior_limb_of_internal_capsule_R | ALIC_R | 18 | 2 | 12 |
| Anterior_limb_of_internal_capsule_L | ALIC_L | -22 | 8 | 16 |
| Posterior_limb_of_internal_capsule_R | PLIC_R | 26 | -20 | 10 |
| Posterior_limb_of_internal_capsule_L | PLIC_L | -18 | -16 | -2 |
| Retrolenticular_part_of_internal_capsule_R | RLIC_R | 30 | -26 | 4 |
| Retrolenticular_part_of_internal_capsule_L | RLIC_L | -34 | -36 | 6 |
| Anterior_corona_radiata_R | ACR_R | 24 | 30 | 8 |
| Anterior_corona_radiata_L | ACR_L | -22 | 28 | 12 |
| Superior_corona_radiata_R | SCR_R | 18 | 6 | 28 |
| Superior_corona_radiata_L | SCR_L | -18 | -16 | 40 |
| Posterior_corona_radiata_R | PCR_R | 26 | -48 | 26 |
| Posterior_corona_radiata_L | PCR_L | -26 | -40 | 26 |
| Posterior_thalamic_radiation_R | PTR_R | 36 | -56 | 4 |
| Posterior_thalamic_radiation_L | PTR_L | -34 | -54 | 2 |
| Sagittal_stratum_R | SS_R | 40 | -28 | -10 |
| Sagittal_stratum_L | SS_L | -40 | -28 | -10 |
| External_capsule_R | EC_R | 34 | -6 | -10 |
| External_capsule_L | EC_L | -32 | 6 | -6 |
| Cingulum_(cingulate_gyrus)_R | CgC_R | 8 | -26 | 34 |
| Cingulum_(cingulate_gyrus)_L | CgC_L | -8 | -26 | 34 |
| Cingulum_(hippocampus)_R | CgH_R | 20 | -40 | -2 |
| Cingulum_(hippocampus)_L | CgH_L | -18 | -40 | -4 |
| Fornix cres or stria terminalis_R | FX/ST_R | 28 | -24 | -4 |
| Fornix cres or stria terminalis _L | FX/ST_L | -28 | -28 | -2 |
| Anterior_Superior_longitudinal_fasciculus_R | ASLF_R | 34 | 4 | 24 |
| Middle_Superior_longitudinal_fasciculus_R | MSLF_R | 34 | -22 | 34 |
| Posterior_Superior_longitudinal_fasciculus_R | PSLF_R | 40 | -46 | 8 |
| Anterior_Superior_longitudinal_fasciculus_L | ASLF_L | -34 | 6 | 22 |
| Middle_Superior_longitudinal_fasciculus_L | MSLF_L | -36 | -34 | 32 |
| Posterior_Superior_longitudinal_fasciculus_L | PSLF_L | -32 | -44 | 30 |
| Superior_fronto-occipital_fasciculus_R | SFO_R | 20 | 0 | 24 |
| Superior_fronto-occipital_fasciculus_L | SFO_L | -22 | 6 | 22 |
| Uncinate_fasciculus_R | UNC_R | 34 | -2 | -14 |
| Uncinate_fasciculus_L | UNC_L | -36 | -2 | -18 |
| Tapetum_R | TAP_R | 30 | -46 | 16 |
| Tapetum_L | TAP_L | -26 | -46 | 18 |

**Table S2.** Nodes for Gray Matter Functional Brain Network Construction

| **Anatomical Regions** | **Abbreviations** | **Local Maxima in MNI coordinates of brain activation** | | |
| --- | --- | --- | --- | --- |
|  |  | **x** | **y** | **z** |
| L. Basal ganglia, globus pallidus | BG_L_6_2 | -22 | -2 | 4 |
| L. Basal ganglia, ventromedial putamen | BG_L_6_4 | -23 | 7 | -4 |
| L. Basal ganglia, dorsal caudate | BG_L_6_5 | -14 | 2 | 16 |
| L. Basal ganglia, dorsolateral putamen | BG_L_6_6 | -28 | -5 | 2 |
| R. Basal ganglia, globus pallidus | BG_R_6_2 | 22 | -2 | 3 |
| R. Basal ganglia, ventromedial putamen | BG_R_6_4 | 22 | 8 | -1 |
| R. Basal ganglia, dorsolateral putamen | BG_R_6_6 | 29 | -3 | 1 |
| L. Cingulate gyrus, caudodorsal | CG_L_7_5 | -5 | 7 | 37 |
| R. Cingulate gyrus, pregenual | CG_R_7_3 | 5 | 28 | 27 |
| L. Fusiform gyrus, medioventral | FuG_L_3_2 | -31 | -64 | -14 |
| L. Fusiform gyrus, lateroventral | FuG_L_3_3 | -42 | -51 | -17 |
| R. Fusiform gyrus, medioventral | FuG_R_3_2 | 31 | -62 | -14 |
| R. Fusiform gyrus, lateroventral | FuG_R_3_3 | 43 | -49 | -19 |
| L. Inferior frontal gyrus, dorsal | IFG_L_6_1 | -46 | 13 | 24 |
| L. Inferior frontal gyrus, opercular | IFG_L_6_5 | -39 | 23 | 4 |
| L. Inferior frontal gyrus, ventral | IFG_L_6_6 | -52 | 13 | 6 |
| R. Inferior frontal gyrus, dorsal | IFG_R_6_1 | 45 | 16 | 25 |
| R. Inferior frontal sulcus | IFG_R_6_2 | 48 | 35 | 13 |
| R. Inferior frontal gyrus, caudal | IFG_R_6_3 | 54 | 24 | 12 |
| R. Inferior frontal gyrus, opercular | IFG_R_6_5 | 42 | 22 | 3 |
| R. Inferior frontal gyrus, ventral | IFG_R_6_6 | 54 | 14 | 11 |
| L. Dorsal agranular insula | INS_L_6_3 | -34 | 18 | 1 |
| L. Dorsal granular insula | INS_L_6_5 | -38 | -8 | 8 |
| L. Dorsal dysgranular insula | INS_L_6_6 | -38 | 5 | 5 |
| R. Dorsal agranular insula | INS_R_6_3 | 36 | 18 | 1 |
| R. Dorsal dysgranular insula | INS_R_6_6 | 38 | 5 | 5 |
| L. Inferior parietal lobule, rostrodorsal | IPL_L_6_2 | -38 | -61 | 46 |
| L. Inferior Parietal Lobule, rostrodorsal | IPL_L_6_3 | -51 | -33 | 42 |
| L. Inferior parietal lobule, caudal | IPL_L_6_4 | -56 | -49 | 38 |
| L. Inferior parietal lobule, rostroventral | IPL_L_6_6 | -53 | -31 | 23 |
| R. Inferior parietal lobule, rostrodorsal | IPL_R_6_2 | 39 | -65 | 44 |
| R. Inferior parietal lobule, rostrodorsal | IPL_R_6_3 | 47 | -35 | 45 |
| R. Inferior parietal lobule, caudal | IPL_R_6_4 | 57 | -44 | 38 |
| R. Inferior parietal lobule, rostroventral | IPL_R_6_5 | 53 | -54 | 25 |
| L. Inferior temporal gyrus, extreme lateroventral | ITG_L_7_2 | -51 | -57 | -15 |
| L. Inferior temporal gyrus, ventrolateral | ITG_L_7_5 | -55 | -60 | -6 |
| L. Inferior temporal gyrus, caudolateral | ITG_L_7_6 | -59 | -42 | -16 |
| R. Inferior temporal gyrus, extreme lateroventral | ITG_R_7_2 | 53 | -52 | -18 |
| R. Inferior temporal gyrus, ventrolateral | ITG_R_7_5 | 54 | -57 | -8 |
| R. Inferior temporal gyrus, caudolateral | ITG_R_7_6 | 61 | -40 | -17 |
| L. Middle occipital gyrus | LOcC_L_4_1 | -31 | -89 | 11 |
| L. lateral occipital cortex | LOcC_L_4_2 | -46 | -74 | 3 |

**Table S2** (Continued). Nodes for Gray Matter Functional Brain Network Construction

| **Anatomical Regions** | **Abbreviations** | **Local Maxima in MNI coordinates of brain activation** | | |
| --- | --- | --- | --- | --- |
|  |  | **x** | **y** | **z** |
| L. Occipital polar cortex | LOcC_L_4_3 | -18 | -99 | 2 |
| L. Inferior occipital gyrus | LOcC_L_4_4 | -30 | -88 | -12 |
| L. Middle frontal gyrus, dorsal | MFG_L_7_1 | -27 | 43 | 31 |
| L. Inferior frontal junction | MFG_L_7_2 | -42 | 13 | 36 |
| L. Middle frontal gyrus | MFG_L_7_3 | -28 | 56 | 12 |
| L. Middle frontal gyrus, ventral | MFG_L_7_4 | -41 | 41 | 16 |
| L. Middle frontal gyrus, ventrolateral | MFG_L_7_5 | -33 | 23 | 45 |
| L. Middle frontal gyrus, ventrolateral | MFG_L_7_6 | -32 | 4 | 55 |
| R. Middle frontal gyrus, dorsal | MFG_R_7_1 | 30 | 37 | 36 |
| R. Inferior frontal junction | MFG_R_7_2 | 42 | 11 | 39 |
| R. Middle frontal gyrus | MFG_R_7_3 | 28 | 55 | 17 |
| R. Middle frontal gyrus, ventral | MFG_R_7_4 | 42 | 44 | 14 |
| R. Middle frontal gyrus, ventrolateral | MFG_R_7_5 | 42 | 27 | 39 |
| R. Middle frontal gyrus, ventrolateral | MFG_R_7_6 | 34 | 8 | 54 |
| R. Middle frontal gyrus, lateral | MFG_R_7_7 | 25 | 61 | -4 |
| L. Middle temporal gyrus, caudal | MTG_L_4_1 | -65 | -30 | -12 |
| L. Anterior superior temporal sulcus | MTG_L_4_4 | -58 | -20 | -9 |
| R. Middle temporal gyrus, caudal | MTG_R_4_1 | 65 | -29 | -13 |
| R. Middle temporal gyrus, dorsolateral | MTG_R_4_3 | 60 | -53 | 3 |
| R. Anterior superior temporal sulcus | MTG_R_4_4 | 58 | -16 | -10 |
| L. Orbital gyrus, lateral | OrG_L_6_3 | -23 | 38 | -18 |
| L. Orbital gyrus, lateral | OrG_L_6_6 | -41 | 32 | -9 |
| R. Orbital gyrus, orbital | OrG_R_6_2 | 40 | 39 | -14 |
| R. Orbital gyrus, lateral | OrG_R_6_3 | 23 | 36 | -18 |
| R. Orbital gyrus, lateral | OrG_R_6_6 | 42 | 31 | -9 |
| L. Postcentral gyrus (upper limb, head and face region) | PoG_L_4_1 | -50 | -16 | 43 |
| L. Postcentral gyrus (tongue and larynx region) | PoG_L_4_2 | -56 | -14 | 16 |
| L. Postcentral gyrus | PoG_L_4_3 | -46 | -30 | 50 |
| L. Postcentral gyrus (trunk region) | PoG_L_4_4 | -21 | -35 | 68 |
| R. Postcentral gyrus | PoG_R_4_3 | 48 | -24 | 48 |
| L. Precentral gyrus (head and face region) | PrG_L_6_1 | -49 | -8 | 39 |
| L. Precentral gyrus, caudal dorsolateral | PrG_L_6_2 | -32 | -9 | 58 |
| L. Precentral gyrus (upper limb region) | PrG_L_6_3 | -26 | -25 | 63 |
| L. Precentral gyrus (trunk region) | PrG_L_6_4 | -13 | -20 | 73 |
| L. Precentral gyrus (tongue and larynx region) | PrG_L_6_5 | -52 | 0 | 8 |
| L. Precentral gyrus, caudal ventrolateral | PrG_L_6_6 | -49 | 5 | 30 |
| R. Precentral gyrus, caudal dorsolateral | PrG_R_6_2 | 33 | -7 | 57 |
| R. Precentral gyrus (tongue and larynx region) | PrG_R_6_5 | 54 | 4 | 9 |
| R. Precentral gyrus, caudal ventrolateral | PrG_R_6_6 | 51 | 7 | 30 |
| L. Rostroposterior superior temporal sulcus | pSTS_L_2_1 | -54 | -40 | 4 |
| L. Caudoposterior superior temporal sulcus | pSTS_L_2_2 | -52 | -50 | 11 |

**Table S2** (Continued). Nodes for Gray Matter Functional Brain Network Construction

| **Anatomical Regions** | **Abbreviations** | **Local Maxima in MNI coordinates of brain activation** | | |
| --- | --- | --- | --- | --- |
|  |  | **x** | **y** | **z** |
| R. Rostroposterior superior temporal sulcus | pSTS_R_2_1 | 53 | -37 | 3 |
| R. Caudoposterior superior temporal sulcus | pSTS_R_2_2 | 57 | -40 | 12 |
| R. Lateral superior occipital gyrus | LOcC_R_2_2 | 29 | -75 | 36 |
| R. Middle occipital gyrus | LOcC_R_4_1 | 34 | -86 | 11 |
| R. Lateral occipital cortex | LOcC_R_4_2 | 48 | -70 | -1 |
| R. Occipital polar cortex | LOcC_R_4_3 | 22 | -97 | 4 |
| R. Inferior occipital gyrus | LOcC_R_4_4 | 32 | -85 | -12 |
| L. Superior frontal gyrus, medial | SFG_L_7_1 | -5 | 15 | 54 |
| L. Superior frontal gyrus, dorsolateral | SFG_L_7_4 | -18 | -1 | 65 |
| L. Superior frontal gyrus, medial | SFG_L_7_5 | -6 | -5 | 58 |
| L. Superior frontal gyrus, medial | SFG_L_7_6 | -5 | 36 | 38 |
| R. Superior frontal gyrus, medial | SFG_R_7_1 | 7 | 16 | 54 |
| R. Superior frontal gyrus, dorsolateral | SFG_R_7_4 | 20 | 4 | 64 |
| R. Superior frontal gyrus, medial | SFG_R_7_5 | 7 | -4 | 60 |
| R. Superior frontal gyrus, medial | SFG_R_7_6 | 6 | 38 | 35 |
| L. Superior parietal lobule, lateral | SPL_L_5_3 | -33 | -47 | 50 |
| L. Superior parietal lobule, postcentral | SPL_L_5_4 | -22 | -47 | 65 |
| L. Superior parietal lobule, intraparietal | SPL_L_5_5 | -27 | -59 | 54 |
| R. Superior parietal lobule, lateral | SPL_R_5_3 | 35 | -42 | 54 |
| R. Superior parietal lobule, intraparietal | SPL_R_5_5 | 31 | -54 | 53 |
| L. Superior temporal gyrus | STG_L_6_2 | -54 | -32 | 12 |
| L. Superior temporal gyrus, caudal | STG_L_6_4 | -62 | -33 | 7 |
| L. Thalamus, medial pre-frontal | Tha_L_8_1 | -7 | -12 | 5 |
| L. Thalamus, pre-motor | Tha_L_8_2 | -18 | -13 | 3 |
| L. Thalamus, sensory | Tha_L_8_3 | -18 | -23 | 4 |
| L. Thalamus, posterior parietal | Tha_L_8_5 | -16 | -24 | 6 |
| L. Thalamus, caudal temporal | Tha_L_8_7 | -12 | -22 | 13 |
| L. Thalamus, lateral pre-frontal | Tha_L_8_8 | -11 | -14 | 2 |
| R. Thalamus, medial pre-frontal | Tha_R_8_1 | 7 | -11 | 6 |
| R. Thalamus, pre-motor | Tha_R_8_2 | 12 | -14 | 1 |
| R. Thalamus, lateral pre-frontal | Tha_R_8_8 | 13 | -16 | 7 |

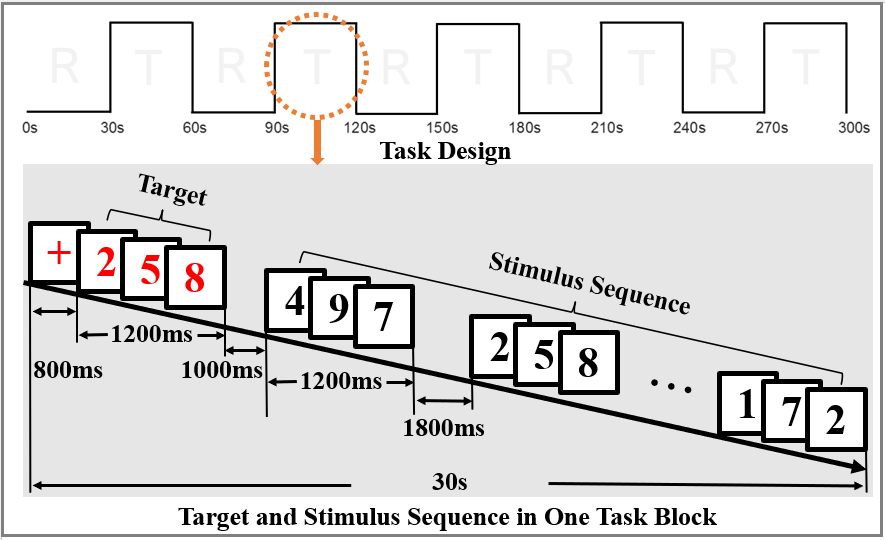

**Figure S1**: Functional MRI experimental task design. (ms: millisecond; s: second)
